# Continuous Titration Based Method For Rapid In-solution Analysis Of Non-covalent Interactions

**DOI:** 10.1101/2024.12.18.629094

**Authors:** Philipp Willmer, Adam C. Hundahl, Rodolphe Marie, Henrik Jensen

## Abstract

Development of new drugs typically involves the identification and validation of molecular inhibitors or promotors of endogenous biological processes. The identification of ligands that can bind the target of interest is typically achieved by screening large libraries of small molecules, using analytical methods that only provide yes/no answers. These methods are only qualitative and often associated with unacceptable amounts of false positives and negatives. Quantitative methods are in general more accurate but time intensive. This is mainly due to repeating measurements of a dilution series in order to generate a titration curve and measure the dissociation constant (*K*_*d*_). In this work, we introduce Continuous Titration Based Spectral Related Intensity Change (cSPRING), that combines Taylor dispersion analysis (TDA) with ratiometric fluorescence detection to measure a *K*_*d*_ in a single experiment. cSPRING is an in-solution method that reduces the sample preparation time 8-fold and requires only nanograms of protein. We show a good agreement of cSPRING with other quantitative methods for three well-known protein-small molecule interactions with binding affinities ranging from the low nanomolar to high micromolar. In addition, we show that cSPRING is able to measure binding affinities in under a minute, highlighting its efficiency and potential for screening applications.

## INTRODUCTION

In biochemical and pharmaceutical research, quantifying the molecular interactions is crucial. Several technologies have been developed to measure these molecular interactions. High throughput screening (HTS) technologies are widely used due to their ability to rapidly evaluate large libraries of compounds. However, these methods often yield only qualitative or semi-quantitative results, typically relying on tracer-based approaches combined with plate readers^[1]^. In contrast, quantitative techniques such as Surface Plasmon Resonance (SPR), Isothermal Titration Calorimetry (ITC), and Bio-Layer Interferometry (BLI) provide comprehensive insights into binding affinities and kinetics. Despite their accuracy, these methods are limited by their medium to low throughput capabilities in the range of tens of minutes to obtain the dissociation constant of a single ligand. Furthermore, each of these methods has inherent limitations. SPR and BLI are both surface-based methods, and so they both require extensive assay development to optimize the surface chemistry and are constrained by buffer compatibility and surface attachment of the analytes, which can affect binding interactions.^[2][3]^ ITC on the other hand is an in-solution technique but necessitates large sample volumes, making it less practical for expensive or rare materials.^[4]^ These constraints, added to their lower throughput, highlight the need for more versatile and efficient analytical techniques.

Taylor dispersion analysis (TDA) is a well-established in-solution technique used to characterize hydrodynamic radii and molecular diffusion.^[5][6]^ In brief, TDA measures the concentration profile of a fluorescent solute as it passes through a capillary under laminar flow. This allows for precise determination of diffusion coefficients and hydrodynamic radii, making TDA ideal for studying biomolecules in solution.^[7][8]^

Flow-induced dispersion analysis (FIDA) evolves from TDA, expanding its application by measuring the concentration profile of a fluorescent target at various ligand concentrations.^[9]^ FIDA has been successfully used to assess molecular stability, binding affinities, and kinetics of non-covalent interactions.^[10][11]^ However, due to its mainly size-change based detection, FIDA is less effective for interactions involving a significantly smaller ligand, such as protein-small molecule binding. In such cases, alternative detection methods are required.

A reliable readout is obtained from the fluorophore’s emission spectrum, as the spectral properties of the fluorescence can be influenced by its environment.^[12][13]^ If the fluorophore is conjugated to a target molecule, a non-covalent interaction with a ligand can alter the microenvironment of the fluorophore and, consequently, its emission spectrum. It has been long known that such subtle changes can be quantified by a ratiometric measurement of intensities in two separate spectral bands in the emission spectrum, which quantifies a spectral related intensity change.^[14][15]^ This technique is inherently self-referenced, making it more robust than a pure fluorescence intensity measurement and has been widely used to characterize non-covalent binding events.^[16][17][18][19]^

We thus combine TDA and FIDA with ratiometric fluorescence detection to present Continuous Titration Based Spectral Related Intensity Change (cSPRING), which offers high throughput quantification of non-covalent interactions to obtain a dissociation constant (*K*_*d*_) and the hydrodynamic radius (*R*_*h*_) of the ligand in seconds to minutes. cSPRING is an in-solution method, which reduces the sample preparation time drastically and requires only nanograms of protein. In brief, the ligand is dispersed under Taylor conditions in a narrow glass capillary.^[5]^ Thereby, it expresses a time dependent, continuous concentration profile as it interacts with a fluorescent target of constant concentration. The change in ratiometric fluorescence is sampled over time, representing a continuous binding curve.

## RESULTS AND DISCUSSION

To demonstrate the working principle of cSPRING, we selected the well characterized protein model system hen egg white lysozyme (HEWL, MW = 14 kDa) which binds the small trisaccharide tri-N-acetyl glucosamine (NAG3, MW = 0.6 kDa) with micromolar affinity.^[20]^ HEWL was conjugated with a Cy5-related fluorescent dye at its lysin groups. This dye is known for having an emission spectrum that is very sensitive towards environmental changes.^[21]^ The dye is excited at *λ*_*ex*_ = 625 nm and the emitted light is collected in two spectral bands from *λ*_*1*_ = 663 – 685 nm and *λ*_*2*_ = 685 – 737 nm. The output of each measurement is the ratiometric fluorescence signal F_*2*_ F_*1*_ ^-1^ over time. A schematic of the setup is shown in Figure S1.

### dSPRING

The interaction of labelled HEWL (referred to as ‘indicator’) and NAG3 (referred to as ‘analyte’) was first studied in a discrete approach to demonstrate the working principle and sensitivity of the ratiometric detection (we will refer to the discrete approach as dSPRING). In this measurement (Figure 1A), the indicator was mixed in a capmix method with different analyte concentrations.^[11]^ In brief, the capillary is filled with analyte, sequentially a small plug (40 nL) of indicator is injected and mobilized with analyte. The indicator arrives after the reference time *t*_*R*_ at the detection window. The fluorescence signal has a Gaussian shape (due to dispersion) and is captured on both detectors.^[5]^ A double Gaussian fit extracts the size related dispersion coefficient σ and the total fluorescence (i.e. the area under the curve) stemming from the indicator and free dye. The ratiometric signal F_2_ F_1_ ^-1^ of the indicator is plotted against the analyte concentration *[A]*. The data is fitted with the model based on Pomorski et al. for ratiometric fluorescence given in Equation (1).^[17]^ Here, *λ*_*b*_ and *λ*_*u*_ are the ratiometric values for fully bound and unbound indicator, respectively. The factor s = F _u1_ F _b1_ ^-1^ relates the intensity of unbound and fully bound indicator in the first spectral band.

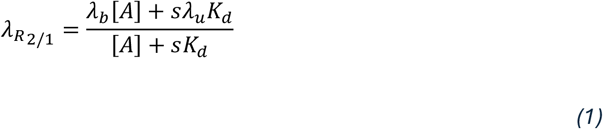

**Figure 1.**
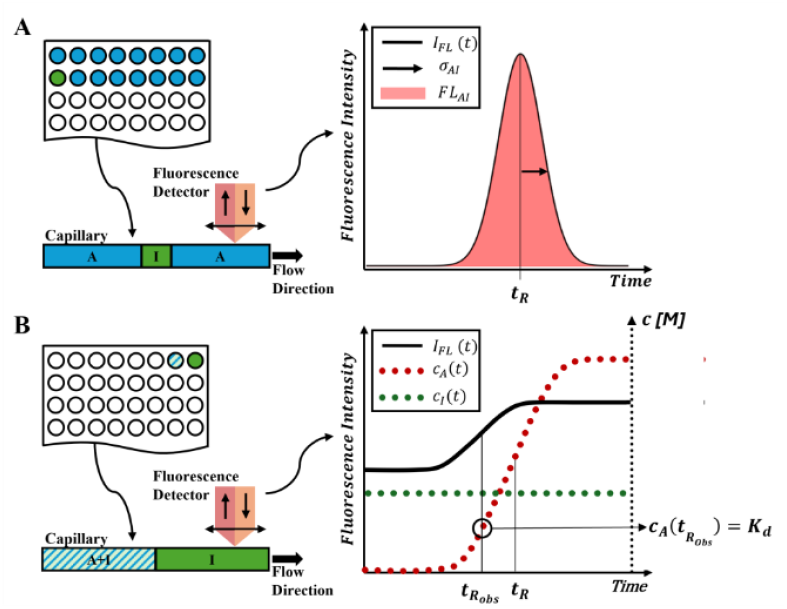
Experimental setup of dSPRING and cSPRING. A: dSPRING - Well-plate with 2-fold dilution series of analyte (blue) and a single indicator well (green). Capmix experiment yields Gaussian-shaped signal B: cSPRING setup – Well plate with 2 samples containing the indicator (green) and the indicator mixed with the analyte (striped), respectively. The indicator fills the capillary and is mobilized with the indicator-analyte mix. Ideally, the indicator concentration (green dots) is constant. The analyte concentration (red dots) can be described analytically by Taylor Dispersion.^[5]^ The inflection point of the fluorescent signal (black solid) indicates where half the indicator is bound and is mapped to a specific analyte concentration which equals the K_d_.

The data with the fitted binding curve and the obtained *K*_*d*_ = 7.2 ± 0.7 µM is shown in Figure 2A. Error bars represent the standard error on the mean of triplicate measurements. The result agrees very well with previously reported *K*_*d*_ values of around 10 µM for this binding system.^[20][22]^

**Figure 2.**
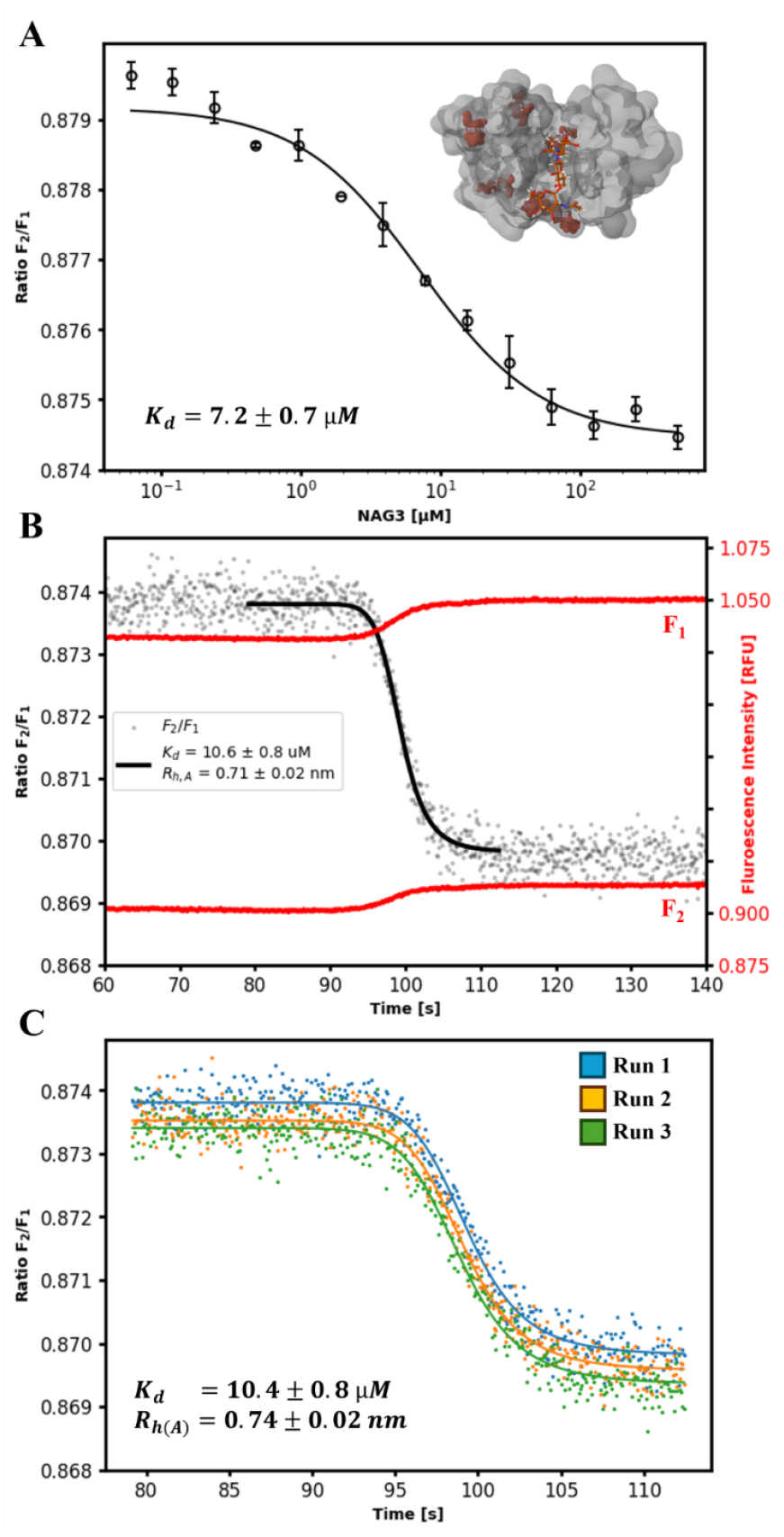
HEWL-NAG3 interaction. A: Discrete 15-point binding curve with 2-fold dilution, start concentration of 500 µM. The data points are the mean of triplicates and error bars represent the standard error on the mean. The top right shows a render of HEWL (grey) - NAG3 (orange) interaction, lysin groups marked in red.^[23][24][25]^ B: cSPRING experimental data, recorded fluorescence signal (red). The ratiometric signal (black scatter) can be clearly resolved. Fit (solid black) yields K_d_ and R_h,A_. C: Triplicate of cSPRING with excellent reproducibility and agreement with previously reported K_d_ values.^[20][22]^

### cSPRING

dSPRING confirms that ratiometric detection can effectively identify binding events. However, it still requires multiple measurements to establish a titration curve. To reduce labor-intensive sample preparation time, reduce uncertainties and accelerate the time-to-result, we introduce a continuous titration-based method (cSPRING). The experimental setup is shown in Figure 1B. Two samples are prepared with the two extreme states of fully bound (indicator mixed with high concentration of analyte, striped in Figure 1B) and unbound indicator (indicator mixed with buffer, green in Figure 1B). The capillary is filled with unbound indicator and mobilized with fully bound indicator. The graph in Figure 1B shows the theoretical concentration profiles (dotted lines) in the capillary and the observed fluorescence signal of a single spectral band (solid black line) as function of time. By design, the total indicator concentration is constant. On the other hand, the analyte is dispersed under Taylor conditions, and its concentration profile can be described analytically according to Equation (2). ^[5][6]^ Here, the equation is expressed in terms of the total analyte concentration *c*_*A*,*0*_, the dispersion coefficient *σ*_*A*_ and the reference time *t*_*R*_, which is the time needed for the interface of the two sample zones to reach the detection window.

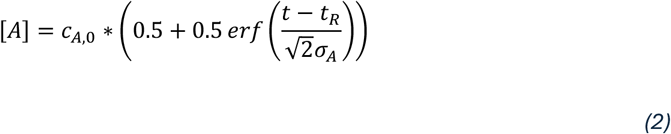

Effectively, Taylor dispersion creates a continuous dilution series of the analyte in a constant indicator concentration, achieving what is normally obtained through time-consuming pipetting. To simplify the concept, we first look at the signal obtained in a single spectral band. Here, the inflection point of the observed fluorescence signal represents where half the indicator is bound. The corresponding time point *t*_*R*,*obs*_ is mapped to the analytically determined concentration profile of the analyte. This yields the *K*_*d*_ value of the interaction. To enhance robustness, cSPRING is based on a ratiometric fluorescence read-out. Unlike single-band fluorescence, which can be affected by experimental artifacts like uneven indicator concentrations, the ratiometric approach mitigates these issues, leading to better accuracy and reliability. Theoretical derivation, simulations and experimental data supporting this method are provided in the Supporting Information.

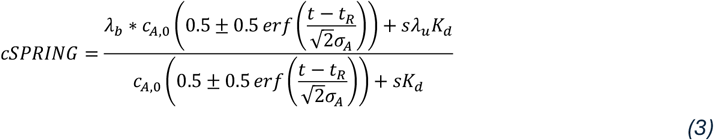

The analytical expression for cSPRING is given in Equation (3). Here, the dispersion coefficient *σ*_*A*_ determines the slope of the ratiometric fluorescence signal and is a fitting parameter. If *σ*_*A*_ is known, it can also be input as a fixed parameter, yielding higher accuracy on the *K*_*d*_ which remains the only unknown parameter. In general, it is assumed that the interaction has fast kinetics and thus, the interaction is always at equilibrium. To ensure that, the experiment can be adjusted by decreasing the flow velocity in the capillary and thus, increasing the time of reaction. Furthermore, the model assumes that the concentration of unbound analyte is approximated with the total concentration of analyte. This assumption is valid for indicator concentrations smaller than the *K*_*d*_ which can be controlled in the experimental setup.

Figure 2B shows the two recorded fluorescence intensities for an indicator concentration of c0,I = 10 nM (red) and the ratiometric signal formed by dividing the two signals (black scatter). The ratiometric signal is fitted with Equation (3), yielding the K_d_ and the dispersion coefficient σ_A_ (black curve in Figure 2B), which is directly related to hydrodynamic radius of the analyte R_h,A_ (Supporting Information). The obtained size of the analyte can give valuable structural information which are otherwise difficult to access. Figure 2C highlights the excellent consistency between a triplicate of the measurement with an average K_d_ = 10.4 ± 0.8 µM and R_h,A_ = 0.74 ± 0.02 nm. The values represent the mean of the three measurements with the error being the standard error on the mean. The K_d_ is similar to the discrete method and also in excellent agreement with previously reported dissociation constants.^[20][22]^ The obtained size of NAG3 is realistic for its molecular weight of MW_NAG3_ = 627 Da.

The cSPRING signal reveals a relative change in the ratio of 0.5 % between the fully bound and unbound state, which is the same as for the discrete method. Also, the absolute values of the ratio are in excellent accordance with each other. The great sensitivity can be explained by the protein structure of HEWL shown in Figure 2A.^[23][24][25]^ HEWL has six lysin residues which are targeted in the labeling reaction. Three of them are in close proximity to the binding cleft (Lys33, Lys96, Lys97), where the greatest micro-environmental alternation is expected, ultimately resulting in a change of the ratiometric signal.

Hereby, cSPRING utilizes only nanograms of indicator (Table S1), optimizing resource use while maintaining high sensitivity. This is particularly interesting where samples are precious or not available in large quantities. Furthermore, cSPRING excels in the drastically improved time-to-result. Since the method uses only two samples, the preparation time is reduced 8-fold. In addition, this makes space in a 96-well plate for quantitatively screening large libraries of small molecules. For example, two 96 well plates should be adequate for more than 150 *K*_*d*_, which using a discrete setup would require more than 1500 wells and extensive sample preparation and comparable large sample amounts.

### Inhibitor screening

To further assess the screening capabilities of cSPRING and its performance for stronger binding interactions, we used bovine carbonic anhydrase II (bCAII) as a model system. This enzyme was selected due to its well-studied interaction with numerous small molecule inhibitors and its extensive characterization using conventional techniques.^[26]^ bCAII was conjugated with the same Cy5-derived fluorescent dye at its lysine residues.

Three different bCAII inhibitors (acetazolamide, furosemide, 4-carboxybenzenesulfonamide) were chosen with *K*_*d*_ values ranging from 20 nM to 1 µM.^[26]^ Triplicates of the cSPRING with the corresponding fits are shown in Figure 3 A-C. The resolved ratio changes range from 0.4 % to 0.6 %, similar to the HEWL. The extracted *K*_*d*_ values and hydrodynamic radii are shown in the bottom left of each plot. Error bars represent the standard error on the mean from triplicate measurements. To further confirm the results, the same samples have been measured in dSPRING, yielding similar binding affinities (Figure S3). These values are consistent with previously reported dissociation constants as summarized in Table 1. Slight differences can be explained by the different measurement methods and potential batch to batch variation of the samples. Also, the obtained hydrodynamic radii agree well with the molecular weight of the small molecules. For further validation, the sizes of the small molecules were measured in a capmix experiment using their native fluorescence (Table S2). The results are in good agreement, where minor differences can be explained by the high concentration of the small molecule and the different buffer conditions.

**Table 1.** Comparison of dissociation constants obtained by different measurement methods.

| Inhibitor | cSPRING | dSPRING | SPR | ITC | Spectral Shift assay |
| --- | --- | --- | --- | --- | --- |
| acetazolamide | $31 \pm 6$ nM | $26 \pm 2$ nM | 19 nM<sup>[26]</sup> | 97 nM<sup>[27]</sup> | $25 \pm 2$ nM<sup>[28]</sup> |
| furosemide | $1002 \pm 125$ nM | $689 \pm 11$ nM | 513 nM<sup>[26]</sup> | 526 nM<sup>[27]</sup> | $426 \pm 63$ nM<sup>[28]</sup> |
| 4-carboxybenzene-sulfonamide | $1761 \pm 79$ nM | $1269 \pm 44$ nM | 913 nM<sup>[26]</sup> | - | - |

**Figure 3.**
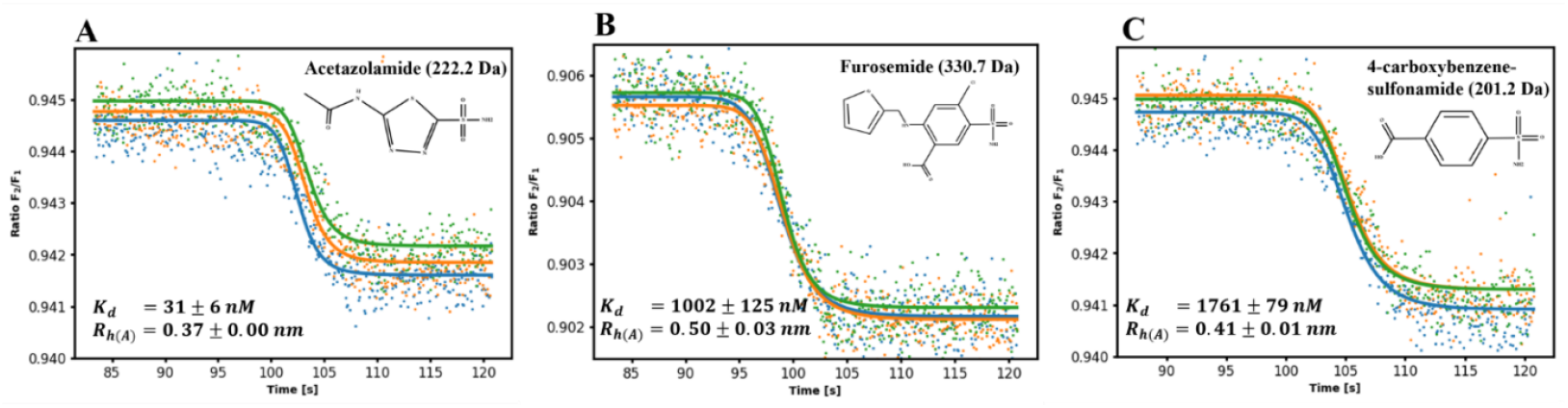
Binding affinities obtained by cSPRING for carbonic anhydrase II with different small molecules inhibitors. A-C: Binding affinity and hydrodynamic radius of the small molecule (structure shown in up-right corner) determined by continuous SRIC for an indicator concentration of 10 nM. Scatter and fitted line of different colors represent the different experiments within the triplicate.

To further highlight the quantitative screening capabilities, cSPRING was performed on a shorter capillary without additional washing steps, reducing the measurement time to 45 seconds for a single *K*_*d*_ value (Figure S4). As a result, triplicates for the three inhibitors were measured in under 7 minutes, consuming only 4.2 µL of indicator solution per inhibitor. Although, the obtained *K*_*d*_ values are less accurate than the ones measured with a longer run time, it allows to discriminate between potential drug candidates on a quantitative basis.

## CONCLUSION

In drug discovery processes, the identification of small molecule ligands that can bind the target of interest is of great interest. High throughput screening of such libraries is often only qualitatively whereas quantitative methods lack speed. In the present work, we propose for the first time cSPRING, a quantitative method which is based on an TDA induced continuous titration and a ratiometric fluorescence readout. cSPRING yields the binding affinity and hydrodynamic radius of the ligand in a single experiment. The method is sensitive towards small molecule binding and requires only one ligand concentration and thus reduces the sample preparation time drastically. This makes it overall suitable to use in automated screening applications with quantitative results. In addition, a single *K*_*d*_ requires only nanograms of protein and is therefore well suited for applications where samples are precious. This positions cSPRING as an ideal solution for accelerating time-intensive pre-screening of non-covalent interactions, both in fundamental research and biopharmaceutical drug development.

## MATERIALS & METHODS

### Equipment

The experiments were performed on a Fida Neo system 640 (Cat-Nr. N002-640, Fida Biosystems ApS). The optical setup of the detection unit is shown schematically in Figure S1. It comprises a light source (LED, *λ*_*peak*_ = 625 nm), an optical arrangement to allow for point detection, two photomultiplier tubes (Hamamatsu) and a dichroic mirror splitting the emitted fluorescent light in two spectral parts. Fused silica capillaries (ID = 75 µm, L = 100 cm) with a length to the detection window of 84 cm (Cat-Nr. 100-001, Fida Biosystems ApS), were used for all experiments. Before each experiment, the capillary is flushed with a coating reagent (Cat-Nr. 310-010, Fida Biosystems ApS).

### Materials and Chemicals

Carbonic anhydrase II from bovine erythrocytes (C2522-1G, Lot-Nr. SLCQ8559) and inhibitors furosemide (F4381, Lot-Nr. MKCM6678), acetazolamide (A6011-10G, Source BCCK0066) and 4-Sulfamoylbenzoic acid (C11804-5G, Source MKCR9857) all purchased from Sigma-Aldrich. Lysozyme from chicken egg white (L6876, Sigma-Aldrich) and trisaccharide tri-N-acetyl glucosamine (OT06497, batch 064971701, Biosynth). Phosphor-buffered saline 1x (PBS) from 10x PBS stock (Lot-Nr. Q01l511, thermo scientific) prepared with deionized water (Direct-Q3, Sigma-Aldrich). TRIS buffer (20 mM, pH 7.8) from 1 M TRIS buffer (15567027, thermos scientific), prepared with deionized water. pH adjusted with 5 M NaOH, from sodium hydroxide pellets (Lot-Nr. SLCC5273, Sigma-Aldrich). Sodium hydroxide 1 M (NaOH) from sodium hydroxide pellets (Lot-Nr. SLCC5273, Sigma-Aldrich)

### Assay buffer

The assay buffer for carbonic anhydrase II is PBS (1x, pH 7.4) with 0.03 % Pluronic F-127 (Millipore, Lot-Nr. 3951203). The assay buffer for lysozyme is TRIS buffer (20 mM, pH 7.8) + 150 mM NaCl (from 5 M NaCl stock solution) with 0.03 % Pluronic F-127 (Millipore, Lot-Nr. 3951203).

### Protein Labeling

Carbonic anhydrase II (bCAII) was dissolved in PBS (1x, pH 7.4) to a final protein concentration of 1 mg/ml. Labelling with an environmentally sensitive dye was performed on the lysine residues using the Fida 1 Protein labelling kit ALC 640 (Cat-Nr. 430-003, Fida Biosystems ApS). A 1 M sodium bicarbonate buffer was added before adding the dye in fourfold molar excess. Incubation time was 30 min in the dark. Excess dye has been removed with the included spin column. The degree of labeling was 0.7 (NanoDrop, Thermo Fisher) and the amount of free dye was below 3 %, measured with a Fida Neo system 640 (Cat-Nr. IN002-640, Fida Biosystems ApS). Labelled bCAII stored at -20 °C.

The hen egg white lysozyme (HEWL) was dissolved in PBS (1x, pH 7.4) to a final protein concentration of approx. 1.4 mg/ml. Labelling with an environmentally sensitive dye was performed on the lysine residues using the Fida 1 Protein labelling kit ALC 640 (Cat-Nr. 430-003, Fida Biosystems ApS). A 1 M sodium bicarbonate buffer was added before adding the dye in fourfold molar excess. Incubation time was 30 min in the dark. Excess dye has been removed with the included spin column. The spin column and resin were rinsed three times with TRIS buffer. Each rinse was performed at 0.8×1000 rpm for 60 seconds. The degree of labeling was 0.9 (NanoDrop, Thermo Fisher) and the amount of free dye was below 5 %, measured with a Fida Neo system 640 (Cat-Nr. IN002-640, Fida Biosystems ApS). Labelled HEWL stored at -20 °C.

### Ligand Preparation

The bCAII-inhibitors furosemide (FS), acetazolamide (AZA) and 4-Sulfamoylbenzoic acid (CBSA) were prepared in DMSO at stock concentrations of 100 mM each at the day of the experiment. The HEWL binding partner trisaccharide tri-N-acetyl glucosamine (NAG3) was prepared at 10 mM in milliQ water. Aliquots were stored at -20 °C and thawed on the day of the experiment.

### dSPRING Experiment

For HEWL-NAG3, the NAG3 stock solution was diluted 20 times in assay buffer to 500 µM. Fifteen-point dilution series (2-fold dilution) were prepared with assay buffer in 96-well plates (Cat-Nr. 220-002, Fida Biosystems ApS), with starting concentration of 500 µM. Labelled HEWL was prepared at a final concentration of 100 nM in assay buffer. The experiments were performed at 25 °C. For bCAII-Inhibitors, the inhibitor stock solutions were diluted a hundred times in PBS to end up with intermediate dilutions of 1 mM. Fifteen-point dilution series (2-fold dilution) were prepared with assay buffer in 96-well plates (Cat-Nr. 220-002, Fida Biosystems ApS), with starting concentrations of 50 µM for FS and CBSA and 5 µM for AZA. Due to very low initial DMSO concentrations (0.05 % for FS, CBSA and 0.005 % for AZA at the first point), no additional DMSO was added to the assay buffer. Labelled bCAII was prepared at a final concentration of 100 nM in an assay buffer. The experiments were performed at 25 °C.

Each measurement required 40 nL of protein and took 5 min analysis time with the following experimental steps: 1) Flush with sodium hydroxide at 3500 mbar for 20 s; 2) Flush with assay buffer at 3500 mbar for 20 s; 3) Flush with coating reagent at 3500 mbar for 20 s. 4) Flush with deionized water at 3500 mbar for 20 s. 5) Fill the capillary with ligand solution at 3500 mbar for 20 s; 6) Inject protein at 50 mbar for 10 s; 7) Mobilize and measure with ligand solution at 400 mbar for 180 s. The signal in both spectral channels was recorded simultaneously for the measurement time during step 7.

### cSPRING Experiment

For HEWL-NAG3, indicator solution was prepared by mixing labelled HEWL and NAG3 with assay buffer to final concentrations of 10 nM and 500 µM, respectively. Analyte solution was prepared by mixing labelled HEWL with assay buffer to final concentration of 10 nM. The experiments were performed at 25 °C. For bCAII-Inhibitors, indicator solutions were prepared by mixing labelled bCAII and bCAII-inhibtiors with assay buffer to final concentrations of 10 nM and 50 µM (5 µM for AZA), respectively. Analyte solution was prepared by mixing labelled bCAII with assay buffer to final concentration of 10 nM.

Each measurement required 8.5 µL of indicator solution and 7 µL of analyte solution and took 4 min analysis time with the following experimental steps: 1) Rinse with sodium hydroxide at 3500 mbar for 25 s. 2) Flush with assay buffer at 3500 mbar for 25 s. 3) Flush with coating reagent at 3500 mbar for 20 s. 4) Flush with deionized water at 3500 mbar for 20 s. 5) Fill the capillary with analyte solution at 3500 mbar for 25 s. 6) Mobilize and measure with indicator solution at 400 mbar for 180 s. The signal in both spectral channels was recorded simultaneously for the measurement time during step 6. The experiments were performed at 25 °C

### Data Processing

For dSPRING experiments, data files for each measurement point were processed in the FIDA analysis software (Fida Software V3.1.1.0, Fida Biosystems ApS). For cSPRING experiments, a custom algorithm was implemented. Detailed description of data analysis in Supporting Information.

## ACKNOWLEDGMENT

Financial support from the Innovation Fund Denmark (grant no. 2052-00002B) is gratefully acknowledged.

## CONFLICT OF INTEREST

The authors declare the following competing financial interest(s): HJ, AH, and PW have commercial interests in FIDA Biosystems ApS. HJ, AH, and PW are employees of FIDA Biosystems ApS.

### Supporting Information

Additional experimental details, data analysis, materials and methods, including schematics of experimental setup (PDF).

## TABLE OF CONTENT

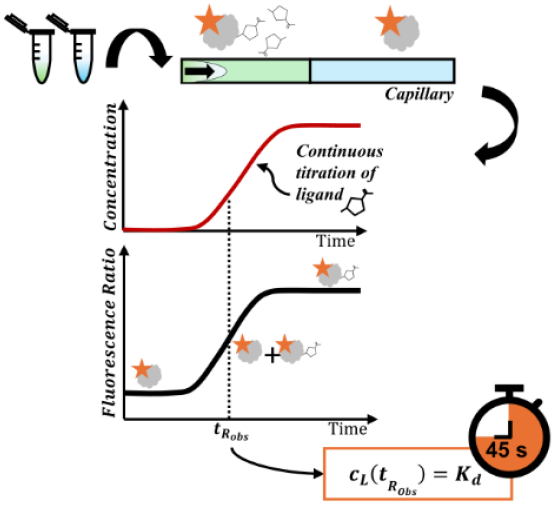

This work presents cSPRING, which combines Taylor dispersion with ratiometric fluorescence detection to measure binding affinities (*K*_*d*_) in a single experiment. This in-solution method reduces sample preparation time 8-fold, requires only nanograms of protein, and provides results in under a minute, demonstrating efficiency for screening protein-small molecule interactions.

