## supplementary file for "Continuous Titration Based Method For Rapid In-solution Analysis Of Non-covalent Interactions"

FIDA Biosystems ApS, 2860 Søborg, Denmark

P. Willmer, R. Marie

Technical University of Denmark, Department of Health Technology, 2800 Kongens-Lyngby, Denmark

**Optical setup**

The used fluorescence detector (Cat-Nr. Det-640N, Fida Biosystems ApS) comprises the optical setup shown in Figure S1. A fibre coupled LED ($\lambda_{ex} = 625\ \mathrm{nm}$) is collimated, passed through a band pass filter (F1, $\lambda = 620/60\ \mathrm{nm}$) and further focused in the flow capillary. The emitted light is collected with the same lens arrangement and passed through a dichroic mirror (DCM 1, $\lambda_{con} = 660\ \mathrm{nm}$) separating the emitted light from the excitation light. The combination of a mirror and a lens couples the emitted light to a fiber, guiding the light to a detection box. Here, the light is collimated again and cleaned up with a band pass filter (F2, $\lambda = 700/75\ \mathrm{nm}$) before being split with a dichroic mirror (DCM 2, $\lambda_{con} = 685\ \mathrm{nm}$) in two portions, covering the two bands as shown in the emission spectrum (shown as insert). The two bands are detected on two individual photomultiplier tubes (PMT 1 and PMT 2). The detector is connected to the software of the supplier (Fida Neo Instrument Software 3.0 Beta).

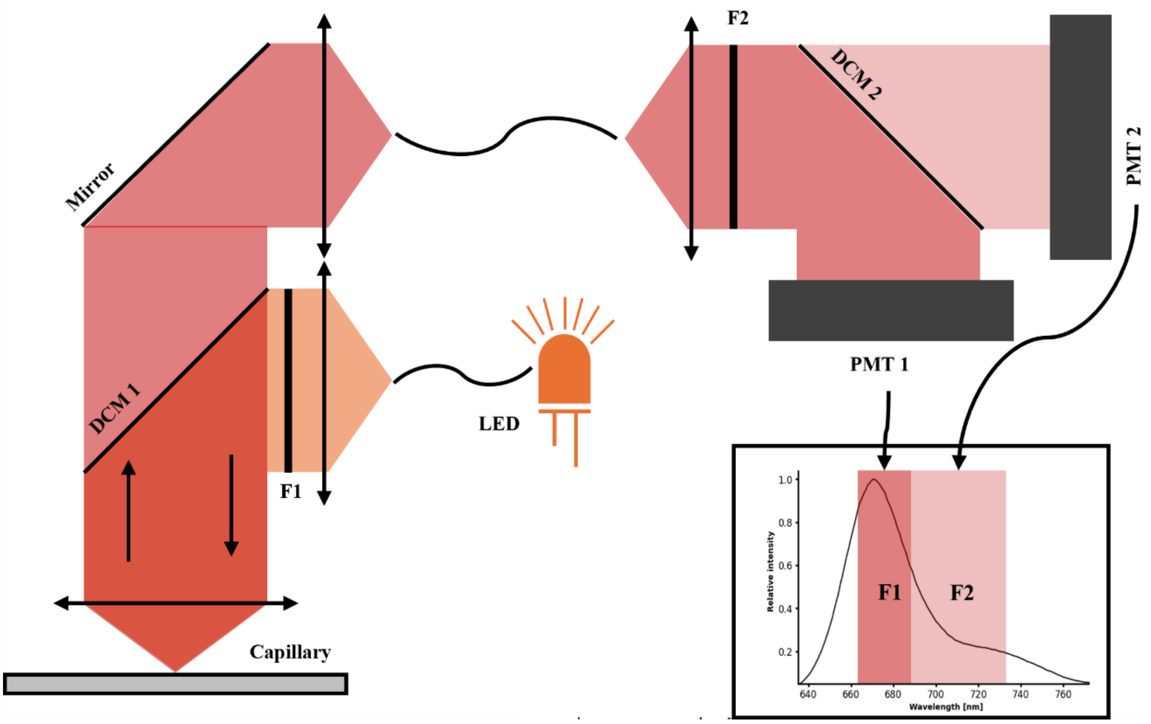


*Figure S1. Optical setup. The light source and the light detectors are coupled with optical fibers to the illumination and detection optics. Excitation and emission share the same optical path. The emission is separated from the illumination by a dichroic mirror (DCM1) and further spectrally filtered by a band pass filter (F2). A second dichroic mirror (DCM2) splits the emitted light in two bands, as seen in the emission spectrum (shown as insert).*

**Protocol for dSPRING data analysis**:

1) Load the data file containing all data points, corresponding to the temporal record of the read-out of PMT1 and PMT2 in the analysis software (Fida Software V3.1.1.0, Fida Biosystems ApS)
2) Obtain the size and the total fluorescence under the curve by performing the automated standard (Gaussian) fit for all fluorescence versus time data:
    a. 2-species fit
    b. 2nd species with fixed $R_h = 0.8\ \mathrm{nm}$ (corresponds to the size of the fluorescent dye)
3) Calculate the ratio from the extracted fluorescent intensities
4) Fit the ratio versus concentration curve using equation (1) and extract the $K_d$

**Protocol for cSPRING data analysis**:

1) Determine the $t_R$ value from the physical constants of the experiment or extract it from a pre-experiment:
    a. Run the fully bound indicator versus buffer in a capmix experiment.
    b. The obtained gaussian can be fitted to extract $t_R$.
2) Block the raw data in blocks of ten data points. Each block is represented by its mean value.
3) Fit the blocked data of the two fluorescence channels with an error function of the form:

$$f(t) = a + b * \operatorname{erf}\left(\frac{t - t_R}{\sqrt{2}\sigma_A}\right)$$

   With the fitting parameters $a, b, t_R$ and $\sigma_A$.

4) Calculate the intensities of fully bound ($F_{b_{1,2}}$) and unbound ($F_{u_{1,2}}$) indicator. (The subscript 1 and 2 indicates the first and the second spectral band, respectively).

$$F_{u_{1,2}} = a_{1,2} - b_{1,2}$$

$$F_{b_{1,2}} = a_{1,2} + b_{1,2}$$

5) Calculate the scaling parameter ($s$) using the fitting parameters:

$$s = \frac{F_{u_1}}{F_{b_1}}$$

6) Calculate the ratiometric parameters $(\lambda_{u,b})$ using the fitting parameters:

$$\lambda_b = \frac{F_{b_2}}{F_{b_1}}$$

$$\lambda_u = \frac{F_{u_2}}{F_{u_1}}$$

7) Calculate the ratiometric signal by dividing the fluorescence signal of detector 2 by detector 1.
8) Fit the cSPRING signal versus time using equation (3) in the region of interest with:
    a. Fixed parameters: $\lambda_u, \lambda_b,\ c_{A,0},\ t_R,\ s$
    b. Fitting parameters: $K_d, \sigma_A$
9) Calculate the hydrodynamic radius $(R_{h,A})$ using the dispersion coefficient $(\sigma_A)$ .

**cSPRING derivation**

The starting point is the ratiometric signal presented by Pomorski et al.[1] where “b” and “u” indicate the fluorescence intensities (F) at fully bound and unbound condition, respectively.

$$\lambda_{R_{2/1}} = \frac{F_{b_2}[A] + F_{u_2}K_d}{F_{b_1}[A] + F_{u_1}K_d}$$

The ratios can be expressed according the respective intensities:

$$\lambda_b = \frac{F_{b_2}}{F_{b_1}} \; and \; \lambda_u = \frac{F_{u_2}}{F_{u_1}}$$

Furthermore, the factor ***s*** is introduced which can be simply readout from the fluorescence intensity of the first spectral band.

$$s = \frac{F_{u_1}}{F_{b_1}}$$

From here, the initial term can be rearranged, starting with inserting $F_{b_1} = F_{u_1}s^{-1}$:

$$\lambda_{R_{2/1}} = \frac{sF_{b_2}[A] + sF_{u_2}K_d}{F_{u_1}[A] + sF_{u_1}K_d}$$

The term is multiplied by $F_{u_1}F_{u_1}^{-1}$:

$$\lambda_{R\,2/1} = \frac{s\frac{F_{b_2}}{F_{u_1}}[A] + s\lambda_u K_d}{[A] + sK_d}$$

Replacing now $F_{u_1} = F_{b_1}s$:

$$\lambda_{R\,2/1} = \frac{\lambda_b[A] + s\lambda_u K_d}{[A] + sK_d}$$

In the discrete SPRING (dSPRING), the above stated equation is fitted to discrete analyte values, yielding the $K_d$.

Plugging in the analytical solution for Taylor dispersion[2],[3] distribution of A, leading to the final expression for the continuous SPRING:

$$cSPRING = \frac{\lambda_b * c_{A,0}\left(0.5 \pm 0.5\,\mathrm{erf}\left(\frac{t - t_R}{\sqrt{2}\sigma_A}\right)\right) + s\lambda_u K_d}{c_{A,0}\left(0.5 \pm 0.5\,\mathrm{erf}\left(\frac{t - t_R}{\sqrt{2}\sigma_A}\right)\right) + sK_d}$$

The equation yields seven parameters. However, the ratios $\lambda_{b,u}$ and the factor $s$ can be directly obtained from the fluorescence signals. The analyte concentration $c_{A,0}$ is given by the experimental design. Lastly, the reference time $t_R$ can be determined analytically for a known setup with a given sample viscosity, temperature and flow velocity or determined in a quick pre-experiment.

Thus, the only two free parameters are the binding affinity $K_d$ and the dispersion coefficient of the analyte $\sigma_A$ which is directly related to its hydrodynamic radius by combining Taylor dispersion and Stokes-Einstein relation:

$$R_{h,A} = \frac{4k_B T\sigma^2}{r_c^2 t_R \pi\eta}$$

Where $T$ is the temperature, $\eta$ is the viscosity and $r_c$ is the radius of the capillary.

**Simulation of the fluorescent signal versus time obtained in a continuous titration experiment (cSPRING)**

An arbitrary indicator-analyte pair is chosen with a total indicator concentration of $c_I = 10\ \mathrm{nM}$ and a total analyte concentration of $c_{0,A}\ =\ 50\ \mu\mathrm{M}$. The reference time is set to $t_R = 105\ \mathrm{s}$. The simulated spectral related intensity change is approximated with 0.5 %, close to experimentally observed change. The signal is normalized to 1 for fully bound indicator.

Figure S1 A shows the fluorescent signal for different $K_d$ values (black curves), and the analytically described concentration profile of the analyte (red). The inflection point of the analyte is at $t_R$. As expected, the curve shifts to the left for lower $K_d$ values, corresponding to a lower analyte concentration needed to fully bind the indicator. For the dotted line, the plateau at 1 is not reached, since the total analyte concentration is too low to fully bind the indicator for the given $K_d$ of 10 µM.

Figure S1 B shows the fluorescent signal for the same arbitrary indicator-analyte pair with an indicator difference $\Delta c_I$ in the two zones. Even concentration mismatches below 5 % skew the signal, making it difficult to analyze. The signals shift to the right. Hence, the observed $t_{R_{obs}}$ value is getting closer to the real $t_R$ value for increasing $\Delta c_I$ since the fluorescence signal reflects the concentration profile of the indicator rather than the cSPRING signal. This artifact is omitted by forming the ratio between the two fluorescent signals. An example is illustrated in Figure S1 C, where the two black curves represent the recorded fluorescent signals, and their ratio is shown in red. The observed $t_{R_{obs}}$ (defined as the midpoint between the plateaus of the fully bound and unbound indicators) closely approximates the actual $t_R = 111\ \mathrm{s}$. Notably, the cSPRING signal reveals the correctly shifted inflection point and eliminates the previously observed bump in the signal.

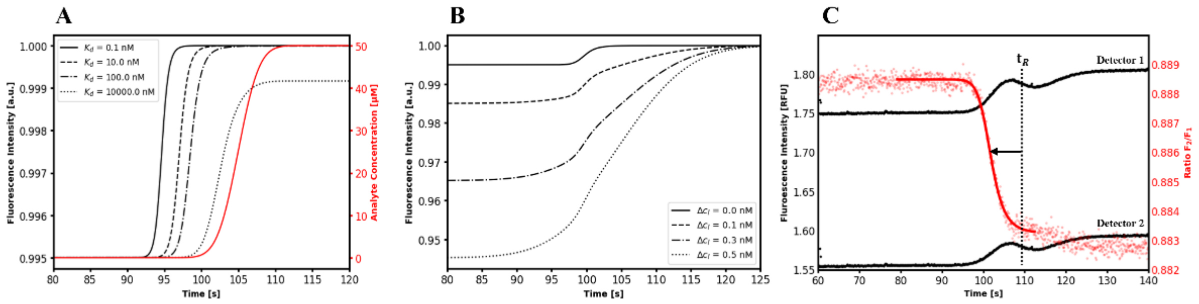


*Figure S2. Simulated data for cSPRING model. A: Fluorescence signals (black) at different $K_d$ values for a given system with same analyte concentration (red). B: Fluorescent signals for indicator mismatches in the two zones. The observed signal is skewed and the observed $t_{R_{obs}}$ value shifts towards the real $t_R$ value. C: Real data of cSPRING signal showing the fluorescence intensity (black) and the ratiometric signal (red). The ratiometric signal reveals the correct slope and shift from the $t_R$ . It eliminates the artifact of the bump which is observable in the fluorescent signal.*

**Calculation of the sample consumption**

Given are stock concentrations of $c_{A,0} = 100\ \mathrm{mM}$ and $c_{I,0} = 1\ \mu\mathrm{M}$ and a minimum volume per well of approx. $V_{min} = 20\ \mu\mathrm{L}$

A 15-point discrete binding curve consumes in a triplicate $41.7\ \mu\mathrm{L}$ per analyte well and a total of $120\ \mathrm{nL}$ indicator. Hence, we select $V_A = 60\ \mu\mathrm{L}$ and $V_I = 25\ \mu\mathrm{L}$. The start concentration for the analyte is $c_A = 50\ \mu\mathrm{M}$ and the indicator concentration is $c_I = 100\ \mathrm{nM}$. This results in a total consumption of stock concentrations of $V_{A_{stock}} = 60\ \mathrm{nL}$ (assuming a volume of $120\ \mu\mathrm{L}$ in the first well for a 2-fold dilution series) and $V_{I_{stock}} = 2.5\ \mu\mathrm{L}$.

In comparison, the continuous binding curve consumes in total for three measurements $22.8\ \mu\mathrm{L}$ of indicator and $18.9\ \mu\mathrm{L}$ of indicator mixed with analyte. For both wells, we select a volume of $V = 50\ \mu\mathrm{L}$. The concentration for the analyte is $c_A = 50\ \mu\mathrm{M}$ and the indicator concentration is $c_I = 10\ \mathrm{nM}$. This results in a total consumption of stock concentrations of $V_{A_{stock}} = 25\ \mathrm{nL}$ and $V_{I_{stock}} = 1\ \mu\mathrm{L}$ ($0.5\ \mu\mathrm{L}$ per well). Overall, the sample consumption is reduced by a factor 2.5 using contiuous titration (cSPRING).

*Table S1. Sample consumption for cSPRING and dSPRING.*

| | Sample | Sample Consumption cSPRING [ng] | Sample Consumption dSPRING [ng] |
|---|---|---|---|
| Protein | Hen Egg White Lysozyme | **14.3** | 36 |
| | Carbonic Anhydrase II | **30** | 75 |
| Small molecules | Tri-N-acetyl glucosamine | **15600** | 37600 |
| | Acetazolamide | **55.5** | 133 |
| | Furosemide | **827** | 1980 |
| | 4-Carboxybenzene sulfonamide | **505** | 1200 |

**Carbonic Anhydrase II + Small molecule inhibitors**

To validate the results of the cSPRING screening of the carbonic anhydrase II, the binding affinities of all three small molecule inhibitors were also measured in the discrete approach. Here we use 15-point binding curves with a 2-fold dilution in PBS (1x, pH 7.4) with starting concentrations of 50 μM for furosemide and 4-carboxzbenzene sulfonamide and 5 μM for acetazolamide. The indicator concentration of the labelled carbonic anhydrase II is 100 nM. The experiment is done in a capmix method with a run-pressure of 400 mbar resulting in an effective indicator concentration of approx. 7.7 nM at the time of detection. The results are shown in Figure S3 A-C. Each data point is the mean of a triplicate where the error bars represent the standard error on the mean. The relative change in the ratiometric signal lies between 0.4 % for acetazolamide and 0.6 % for furosemide, also reflected in the size of the errorbars.

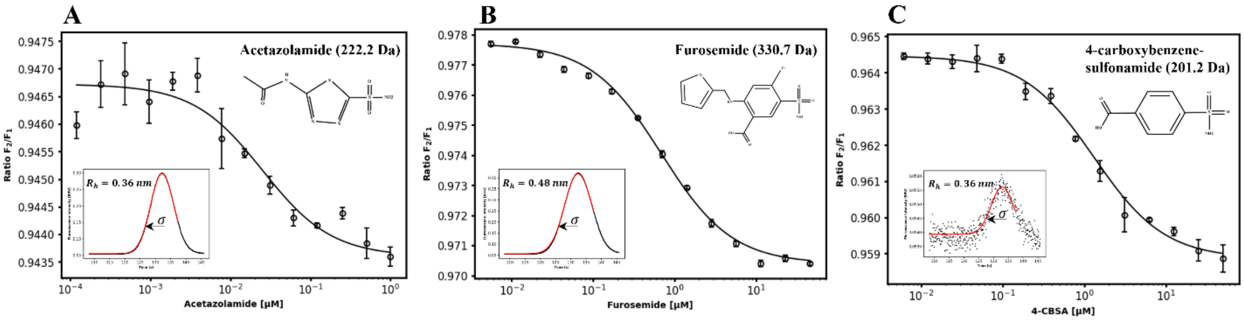


*Figure S3. A-C shows dSPRING data for carbonic anhydrase II with different small molecule inhibitors. The top-right of each figure shows the molecular weight and structure of the small molecule. The bottom-left of each figure shows the FIDA signal obtained from measuring the native fluorescence of the small molecule, exciting them at 280 nm with a UV-fluorescence detector (Cat-Nr. Det-UVN, Fida Biosystems ApS).*

To validate the $R_h$ values obtained via the cSPRING method, the size of each small molecule was measured in a FIDA experiment. Here, we excited the molecules at 280 nm and recorded the fluorescent intensity of their native fluorescence. For all three, the concentration was 100 mM in PBS (1x, pH 7.4) + 10 % DMSO. Examples of the obtained signals are shown in the bottom-left corners in Figure S3 A-C. The hydrodynamic radii were extracted from the width of the peak $\sigma$ (indicated with an arrow), matching very well with the results from cSPRING and the expected sizes, given the molecular weights.

All results are tabulated in Table S2. The error for the $K_d$ is the error on the fit whereas the hydrodynamic radius is given by the mean and the standard error of the mean of a triplicate measurement.

*Table S2. Dissociation constants and hydrodynamic radii obtained for all three small molecule inhibitors from dSPRING and Taylor Dispersion Analysis, respectively.*

| | **acetazolamide** | **furosemide** | **4-carboxybenzene-sulfonamide** |
|---|---|---|---|
| $\boldsymbol{K_d}$ $[\boldsymbol{nM}]$ | $26 \pm 2$ | $689 \pm 11$ | $1269 \pm 44$ |
| $\boldsymbol{R_h}$ $[\boldsymbol{nm}]$ | $0.36 \pm 0.00$ | $0.48 \pm 0.00$ | $0.36 \pm 0.04$ |

**Fast Screening**

To further reduce the measurement time, cSPRING was performed in a shorter capillary with higher pressure and no cleaning steps. A commercially available coated glass capillary (ID = 75 µm, Cat-Nr. 100-002 Fida Biosystems ApS) was shortened to a length of 50 cm with a distance of 34 cm from the inlet to the detection window.

Each measurement required 1.3 µL of indicator solution (bCAII mixed with inhibitor) and 1.9 µL of analyte solution (bCAII) and took 45 s analysis time with the following experimental steps: 1) Fill the capillary with analyte solution at 1500 mbar for 15 s. 2) Mobilize and measure with indicator solution at 500 mbar for 30 s. The experiments were performed at 25 °C.

In summary, triplicates for the three inhibitors were measured in under 7 minutes, consuming only 4.2 µL of indicator solution per inhibitor and a total of 17 µL of analyte solution.

The $K_d$ and $R_{h,A}$ values obtained from triplicate-measurements are displayed in Figure S4 A-C. The result represents the mean of the triplicates, with the error given as the standard error of the mean. Although the results are less accurate than previous measurements, they still provide a quantitative indication of the $K_d$, within agreement of previously reported affinities. Notably, the apparent hydrodynamic radius of the small molecules is larger, which is consistent with the observed deviation in the fitted slope compared to the experimental data. This suggests that the system may have entered a regime where the assumption of fast kinetics and constant equilibrium is no longer valid. The increased hydrodynamic radius indicates that the kinetics is too slow for the system to reach equilibrium in the time it travels to the detection window, resulting in a measured size closer to that of the protein, which is expected under conditions of very slow kinetics.

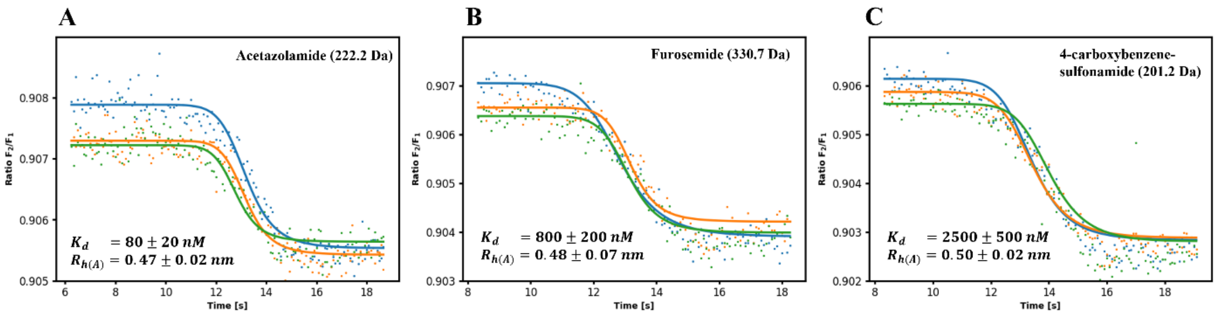


*Figure S4. cSPRING performed in a shorter capillary to decrease the measurement time for a single $K_d$ to 45 seconds. A-C show the cSPRING signals for the three different inhibitors with the fitted $K_d$ and hydrodynamic radius. Although the results are less accurate, the quantitative readout gives valuable insights and is still within agreement of previously reported affinities.*

**Supporting Information – References**

[1] A. Pomorski, T. Kochańczyk, A. Miłoch, A. Kręzel, *Anal Chem* **2013**, *85*, 11479.

[2] U. Sharma, N. J. Gleason, J. D. Carbeck, *Anal Chem* **2005**, *77*, 806.

[3] Geoffrey Ingram Taylor, *Royal Society* **1953**, *219*.
